# Allele Specific Expression Quality Control Fills Critical Gap in Transcriptome Assisted Rare Variant Interpretation

**DOI:** 10.1101/2025.05.30.657086

**Authors:** Kaushik Ram Ganapathy, Eric Song, Daniel Munro, Ali Torkamani, Pejman Mohammadi

## Abstract

Allele-specific expression (ASE) captures the functional impact of genetic variation on transcription, offering a high-resolution view of cis-regulatory effects, but its quality can be diminished by technical, biological, and analysis artifacts. We introduce aseQC, a statistical framework that quantifies sample-level ASE quality in terms of the overall expected extra-binomial variation to exclude uncharacteristically noisy samples in a cohort to improve robustness of downstream analyses. Applying aseQC to a dataset of rare mendelian muscular disorders, successfully identified previously annotated low-quality cases demonstrating clinical genomic utility. When applied to 15,253 samples in extensively quality controlled GTEx project data, aseQC uncovered 563 low-quality samples that exhibit excessive allelic imbalance. We identify these to be associated with specific processing dates but not otherwise described adequately by any other quality control measures and metadata available in GTEx data. We show that these low-quality samples lead to 23.6 and 31.6 -fold increased ASE, and splicing outliers, degrading the performance of transcriptome analysis for rare variant interpretation. In contrast, we did not observe any adverse effect associated with inclusion of these samples in common-variant analysis using quantitative traits loci mapping. By enabling quick and reliable assessment of sample quality, aseQC presents a critical step for identifying subtle quality issues that remain critical for a successful analysis of rare variant effects using transcriptome data.

## Introduction

Allele-specific expression (ASE) quantifies allelic imbalance at the transcript level, providing critical insights into gene regulation and the functional impact of genetic variation in cis [1, 2]. As with any other data, the quality of ASE data is critical to any downstream analysis. Since, ASE integrates signals from DNA and RNA, its quality is ensured using standard DNA and RNA sequencing quality control measures and methods [3-5]. Additionally, tools have been developed to ensure overall matching of the genotypes between the DNA and RNA data used for ASE data generation [6, 7]. Beyond these, ASE data can be distorted by other factors, such as genotyping quality, library preparation, sequencing, and data analysis artifacts, RNA contamination from outside sources and other more subtle sources that together can obscure biologically meaningful ASE signal and limit its usability. To address this gap, we developed aseQC, a statistical method that quantifies deviations from expected biological variance to systematically identify noisy transcriptomes that can compromise downstream analyses. We first validated aseQC in a controlled setting using 40 GEUVADIS RNA-seq samples synthetically perturbed to simulate RNA-seq/DNA-seq mismatches, demonstrating its sensitivity to technical artifacts that distort allelic balance [3]. As a benchmark for clinical usability, we next applied aseQC to a cohort of individuals with inherited rare muscle disorders and successfully recovered all transcriptomes previously flagged as low quality through manual curation [8]. We then scaled the analysis to over 15,000 samples spanning 54 tissues in the GTEx project, where aseQC identified 563 low-quality transcriptomes that had passed existing DNA-seq and RNA-seq quality control pipelines [4]. These samples exhibited inflated rates of ASE and splicing outliers, consistent with pervasive transcriptional noise. Excluding aseQC-flagged samples led to marked improvements in signal fidelity, particularly enhancing the discovery of rare, large-effect regulatory variants. Finally, to assess generalizability across diverse populations, we produce ASE and applied aseQC to an ancestrally diverse cohort of 731 individuals spanning 26 ancestry groups and observed consistent performance across groups [9]. Subsequently, within this dataset, we benchmarked ASE aggregation strategies and found that haplotype-level aggregation consistently improved ASE quality relative to single-SNP approaches, consistent with prior observations, and underscoring the value of phasing for extracting robust regulatory signal from complex transcriptomes [10].

## Results

### Quantifying Extra-Binomial Variation in ASE Data

Population-level allele-specific expression (ASE) data can be conveniently described using the Binomial Logit-Normal (BLN) distribution, parameterized by a mean and variance that capture allelic imbalance and its variability[1, 8]. We extend this framework to model sample-level ASE, where the mean parameter μ represents the average log allelic imbalance, typically reflecting reference bias, and the standard deviation σ captures extra-binomial variability in log allelic imbalance across all genes. However, fitting the BLN model across whole samples can become numerically unstable due to the influence of outlier genes with extreme imbalance and high read counts, which may cause the likelihood function to underflow. To mitigate this, we introduce a uniform-BLN mixture model (Methods, Eq. 4) that enhances numerical stability and enables robust genome-wide inference. (Methods, Eq. 4). The estimated standard deviation (σ) from the BLN component, referred to as the aseQC score, σ(BLN), provides a transcriptome-wide summary of allelic dispersion. To identify samples with globally aberrant ASE, we apply a skew-adjusted boxplot to the distribution of aseQC scores across the cohort, yielding a robust threshold (σ_t_) for detecting outlier samples [11]. We implement this approach in a lightweight pipeline named aseQC which given a cohort estimates the aseQC score for each sample in a cohort and flags outliers using the derived threshold (Fig. S1). aseQC is available as an R package or within a container, supports parallel execution, and runs in seconds per sample (Fig S2 A-B, [12, 13]).

As a first test to validate the effectiveness of aseQC, we simulated genotype–RNA-seq mismatches, which are a well-established source of technical contamination known to distort allele-specific expression (ASE) measurements [6, 7]. Specifically, we introduced synthetic mismatches into RNA-seq data from 40 lymphoblastoid cell line (LCL) samples from the GEUVADIS consortium to create controlled levels of contamination. Applying aseQC to this synthetically perturbed dataset, we observed a strong correlation between the estimated aseQC score, and the degree of mismatch introduced (Fig. 1A–D) [3]). Next, to benchmark the performance of aseQC, we compared it against DROP-MAE QC, a method previously developed for detecting mismatches between genotypes and RNA-seq data leveraging ASE data (Fig. S3A–B, [6]). While both approaches successfully identified contaminated samples, aseQC exhibited enhanced responsiveness across a range of contamination levels and was particularly sensitive to low contamination (Fig. S3C-E).

**Figure 1:**
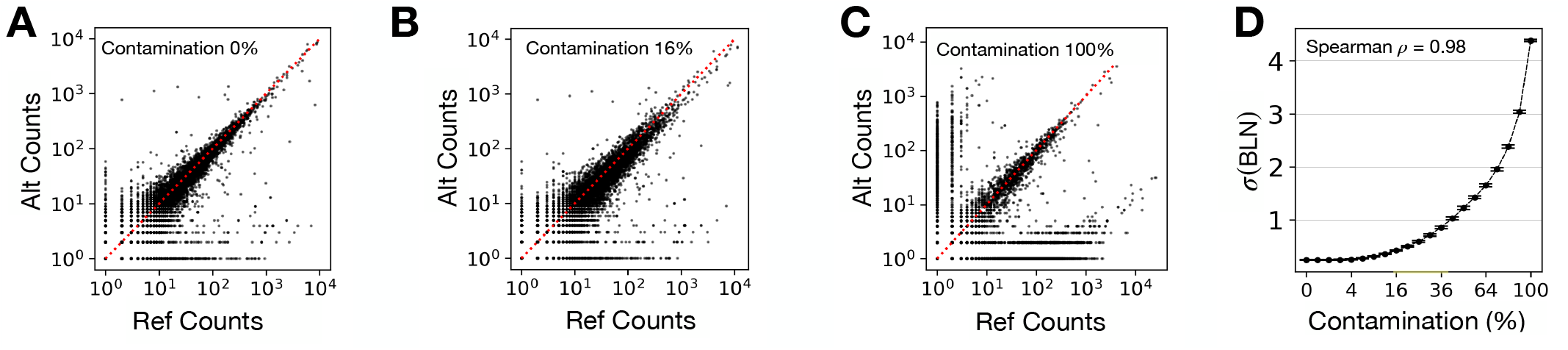
Quantifying Quality in Sample-Level ASE Data Using BLN Distribution. **(A-C)** Allele-specific expression (ASE) distributions for the GEUVADIS sample HG00139 at contamination levels of 0% (A), 16% (B), and 100% (C). **(D)** Relationship between synthetic contamination levels and the fitted BLN standard-deviation (σ) across 40 GEUVADIS samples, with error bars representing the 95% confidence interval (CI) of the mean σ at each level.

### aseQC Identifies Low Quality ASE Samples in a Clinical Rare Disease Cohort

Next, we applied the aseQC pipeline to a clinically relevant cohort of 70 previously analyzed muscular dystrophy cases to validate if it captures samples previously annotated as low quality by manual inspection [8]. In the original study, three samples (N12, N17, and N39) had been excluded due to aberrant ASE profiles, characterized by an unusually high number of genes exhibiting significant allelic imbalance [8]. When aseQC was applied to the full cohort, four samples failed the test, namely, D9, D14, N12, and N39. (Fig. S4, Table S1). Of these, N12 and N39 correspond to previously excluded poor-quality samples, while D14 was excluded but due to low sequencing coverage. The remaining sample, D9, was not excluded in the original analysis and shows an aseQC score marginally exceeding the threshold, suggesting it may represent a borderline case. In contrast, sample N17—previously excluded as low-quality was not flagged by aseQC suggesting that certain forms of aberrant ASE may not produce elevated global allelic dispersion detectable by the model. Collectively, these demonstrates that aseQC not only recapitulates expert-based quality exclusions but also extends sensitivity to additional potentially problematic samples, thereby facilitating automated, reproducible detection of low-quality ASE data in rare-disease settings.

### Applying aseQC to GTEx Data Identifies 563 Failed Samples Across Tissues

We applied aseQC to 15,253 samples across 52 tissues and 2 cell-lines, encompassing 838 individuals from the Genotype-Tissue Expression (GTEx) consortium v8 (Table S2). The pipeline was run on each tissue type independently. We found a wide variability in aseQC scores and thresholds across the tissue types with whole blood samples at the highest threshold (0.49), and fibroblasts at the lowest (0.26) (Fig 2A). Across all tissues, 563 samples (3.69%) did not pass the aseQC test. In tissues with over 25 samples, adrenal gland had the fewest proportion of rejected samples (0.43%), whereas whole blood had the most (7.46%) (Fig 2B). Overall, the median percentage of samples rejected was 3.68% per tissue (Fig 2B). To determine whether aseQC-failed samples are captured by existing genotype-RNA mismatch frameworks, we applied the DROP-MAE QC module [6]. We applied DROP-MAE QC to all 563 aseQC-failed samples across tissues, and found that only 2 were flagged (Fig. S5, Table S3, [6]). This result is consistent with extant quality control procedures in GTEx analysis pipeline, which include verifying genotype–RNA concordance as part of the eQTL mapping workflow [4].

**Figure 2:**
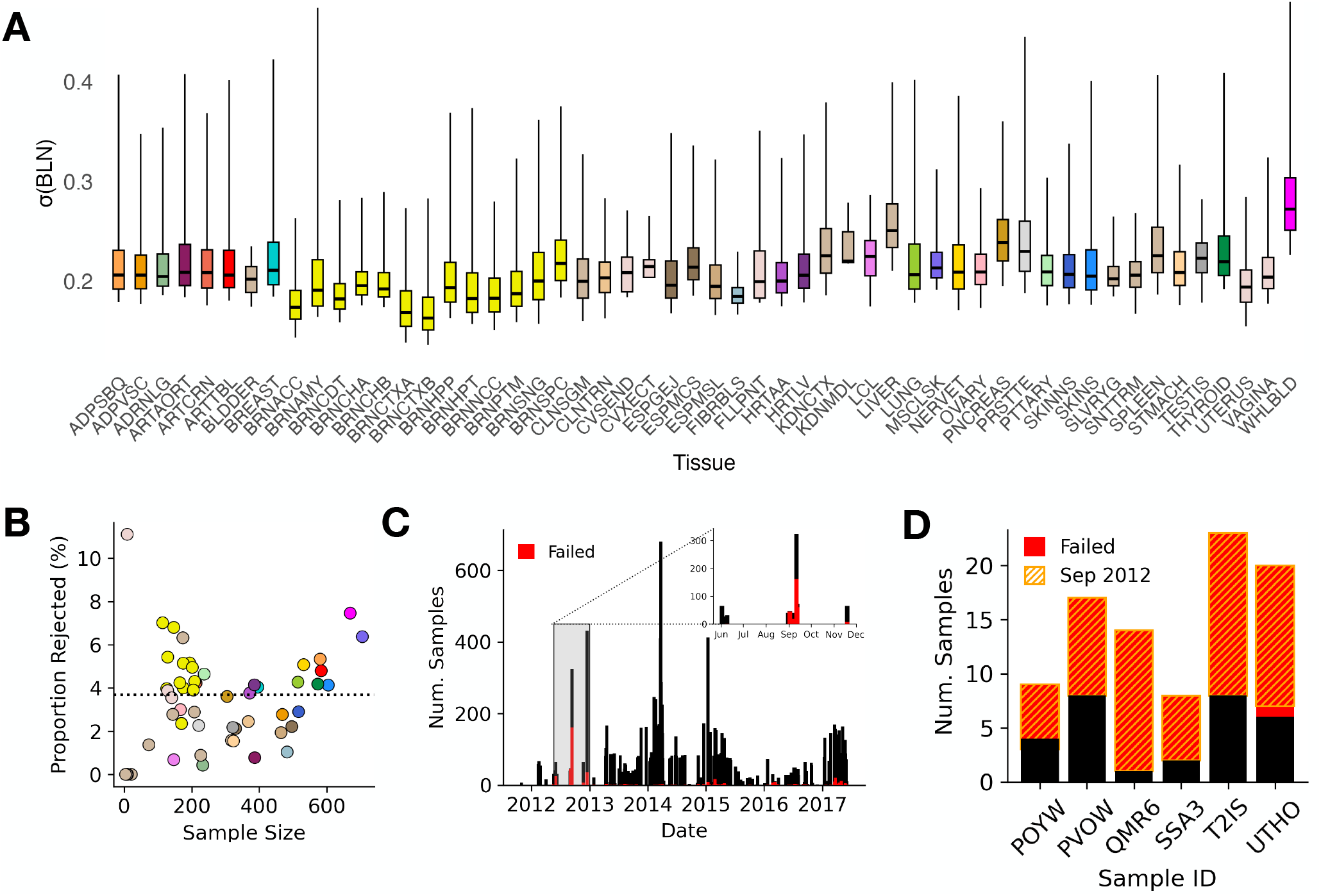
Applying aseQC Across GTEx Tissues Reveals Temporal Trends in ASE Quality. **(A)** Skew-adjusted boxplots of σ(BLN) for 52 tissues and 2 cell lines in GTEx. **(B)** Percentage of rejected samples per tissue plotted against tissue sample size, with points color-coded by tissue. **(C)** Trends in aseQC outcomes over time, highlighting a surge in failures in September 2012. **(D)** Donors with more failed samples than passed, all of whom had a significant portion of their samples processed in September 2012.

### Sample Attributes Associated With aseQC Score in GTEx

Next, we trained a random forest model to assess whether aseQC scores could be reliably predicted from sample-level attributes, including RNA integrity number (RIN), sequencing depth, and alignment metrics such as mapping rate and duplication rate. Model training was performed with cross-validation and hyperparameter tuning to optimize predictive performance (Fig. S6A). Predicted scores from the training set were used to construct skew-adjusted boxplots, from which tissue-specific thresholds for outlier classification were derived. Performance on an independent test set was evaluated using standard benchmarking metrics, including precision, recall, F1-score and the Matthews correlation coefficient (MCC, r score), a classification equivalent of the Pearson correlation coefficient (Fig S6G., [14, 15]). We categorized metadata—drawn from published GTEx annotations—into three main groups: (1) RNA-seq quality control metrics generated by RNA-SeQC, (2) temporal features such as batch processing dates, and (3) pre-sequencing variables including tissue type, RNA integrity number (RIN), and ischemic time. (Table S4). To evaluate the contribution of different feature sets, we first trained a baseline model using only pre-sequencing attributes, which yielded an r-score of 0.09—indicative of near-random performance (Fig. S6B, F). Incorporating RNA-seq quality metrics alongside pre-sequencing attributes substantially improved predictive power, increasing the r score to 0.46 (Fig. S6C, F). Notably, a model combining pre-sequencing attributes with only two temporal features—batch processing dates (SMGEBTCHD and SMNABTCHD)—achieved an even higher r-score of 0.55 (Fig. S6D, F), suggesting that temporal metadata alone provides greater predictive value than 48 RNA-seq QC features. Finally, the full model incorporating all available features attained the highest r score of 0.57 (Fig. S6E-F). Similar trends were observed across other performance metrics, including precision, recall, and F1-score (Fig. S6G). While this reflects a weakly moderate predictive accuracy, it highlights that the aseQC score cannot be fully explained by sample-level metadata alone. To further elucidate which sample attributes most strongly influenced ASE quality, we applied SHAP (SHapley Additive exPlanations) to the trained random forest model. SHAP provides a theoretically grounded approach for interpreting machine learning predictions by quantifying the marginal contribution of each feature to the model’s output. This allowed us to disentangle the relative importance and directionality of each predictor contributing to aseQC score variation across samples. Among all features examined, the most influential predictor was the expression or genotype batch date (SMGEBTCHD), highlighting the outsized impact of temporal batch effects (Fig. S6D, H). Other features positively associated with aseQC score included increased RNA integrity (SMRIN), higher mapping rate (SME1MPRT), more accurate strand-specific alignment (SME1PCTS), and reduced rRNA contamination (SMRRNART). These attributes are consistent with known determinants of RNA-seq data quality and further validate the biological relevance of the aseQC score.

To further elucidate which sample attributes most strongly influenced the aseQC score, we applied SHAP (SHapley Additive exPlanations) to the trained random forest model (Fig S7A-H, [16]). SHAP provides a theoretically grounded approach for interpreting machine learning predictions by quantifying the marginal contribution of each feature to the model’s output. This allowed us to disentangle the relative importance of each predictor contributing to aseQC score variation across samples. Among all features examined, the most influential predictor was the expression or genotype batch date (SMGEBTCHD), highlighting the outsized impact of temporal batch effects (Fig. S7D, H). Other features positively associated with higher aseQC score included increased RNA integrity (SMRIN), higher mapping rate (SME1MPRT), more accurate strand-specific alignment (SME1PCTS), and reduced rRNA contamination (SMRRNART). These attributes are consistent with known determinants of RNA-seq data quality and further validate the biological relevance of the aseQC score.

Finally, we evaluated whether samples failing aseQC were disproportionately concentrated in specific individuals, which could indicate donor-specific technical or biological confounders. Of the 838 individuals in GTEx v8, the majority (n = 598; 71.36%) had no samples that failed aseQC across any tissue (Table S2). Among the remaining 240 individuals, the median number of failed samples per-individual was one. However, a small subset (n = 6; 0.71%) exhibited more failed than passed samples (Fig. 2D). Closer examination revealed that these cases aligned with distinct processing windows (see following section), implicating temporally localized batch effects—rather than inherent donor-related issues—as the primary driver of these failures. Taken together, these findings suggest that failure in aseQC is predominantly driven by batch-specific effects, which are not fully captured by available metadata or conventional RNA-seq quality control metrics.

### aseQC Failures are Clustered in Specific Sample Processing Periods

To investigate the impact of sample processing dates (SMGEBTCHD) on ASE quality, we plotted the number of samples that passed or failed the aseQC test over time (Fig 2C, S8A-D). We observed a significant spike in failures in September 2012 and March 2016. 283 out of 485 samples in September 2012 (58.3%) and 12 out of the 14 samples (86.7%) in March 2016 failed the aseQC test. (Fig S8B). Collectively, samples failing during these spikes accounted for over half of all failed samples across the entire GTEx dataset (Fig S8C). When stratified by individual IDs, all six individuals with more failed than passed samples were found to have most of their samples processed during the September 2012 spike, with nearly all these samples resulting in failure (Fig 2D). The constancy in genotyping across individuals, suggests that the observed variations in aseQC score across GTEx are likely attributable to the processing date of the RNA-sequencing (transcriptomic) data.

### Samples Failing aseQC Show Broader Distortion of Transcriptional Signals Beyond ASE

To determine whether aseQC-failed samples exhibit broad transcriptomic disruption, we first performed principal component analysis (PCA) on log-transformed gene expression profiles across 49 GTEx tissues with at least 25 samples (Table S5). Projection into the top four principal components—collectively explaining over 50% of variance—revealed that aseQC-failed samples were, on median, 1.49-fold farther from the tissue-specific expression centroid than aseQC-passing samples (Wilcoxon signed-rank test; p: 3.35 × 10^−9^), indicating deviations in global expression structure in these samples (Figure S9C).

To further assess the impact of aseQC failures across transcriptional phenotypes, we identified genes in each individual that are outlier for gene expression, splicing, and allele-specific expression (ASE) across GTEx. We applied OutSingle for gene expression outliers, FRASER2 for splicing outliers, and ANEVA-DOT for ASE outliers, focusing on 49 GTEx tissues with at least 25 samples [8, 17, 18]. For each transcriptional phenotype, we computed the mean gene-level outlier rate, stratified by aseQC status. aseQC-failed samples exhibited markedly elevated outlier rates: on average, they harbored 23.58-fold more ASE outliers, 31.60-fold more splicing outliers, and 1.42-fold more gene expression outliers compared to aseQC-passing samples (Fig. 3A). To quantify the statistical significance of this inflation across tissues, we performed Wilcoxon rank-sum tests comparing outlier burdens between aseQC-passing and -failing samples. Significant inflation was observed in 4 tissues for gene expression outliers, 47 tissues for splicing outliers, and 48 tissues for ASE outliers (Fig. S10A–AW). These findings underscore the distortion of transcriptional signals in samples that fail aseQC which can potentially impact transcriptome analyses that do not directly concern ASE data.

**Figure 3:**
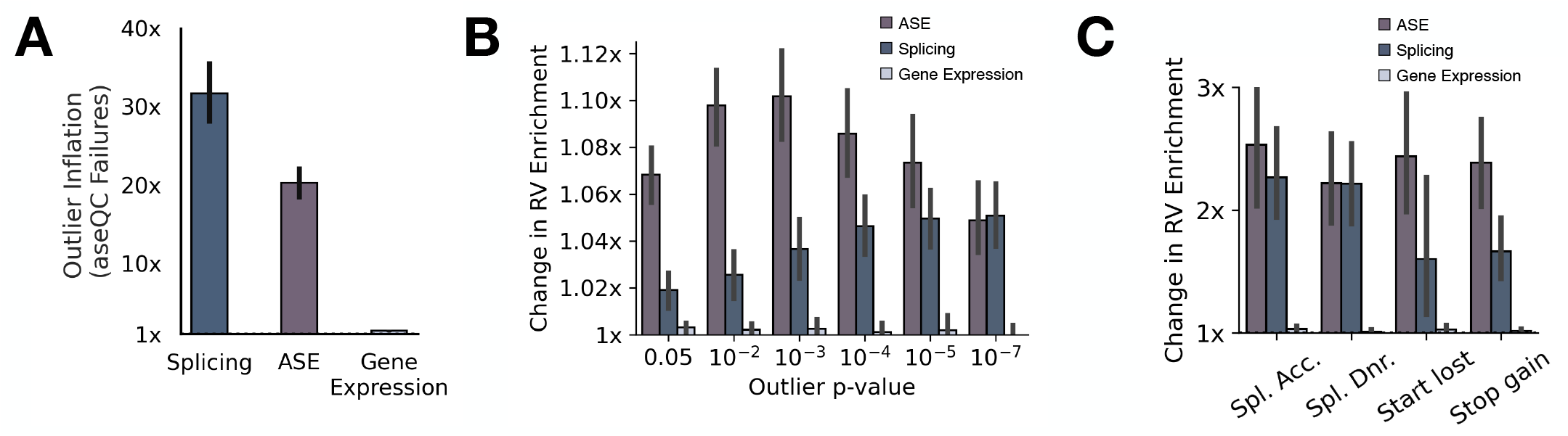
aseQC Failures Alter Outlier Rates and Rare Variant Enrichment Across Transcriptional Phenotypes. **(A)** Mean outlier inflation, defined as the ratio of outlier gene proportions in aseQC failures to passes, across transcriptional phenotypes. Error bars represent the 95% confidence interval (CI) of the mean across tissues with sufficient sample sizes (n > 25). **(B)** Mean change in rare variant enrichment stratified by outlier significance, with 95% CI error bars across eligible tissues. **(C)** Mean change in rare variant enrichment across high-impact VEP classes, with 95% CI error bars shown for annotations present in at least 45 of 49 tissues.

### Removing Samples That Fail aseQC Improves Enrichment for Rare Variants Affecting Allelic Imbalance and Alternative Splicing

We next evaluated the impact of aseQC-based filtering on downstream regulatory variant analyses, focusing on both common and rare variant associations across transcriptional phenotypes. Expression quantitative trait loci (eQTLs) and splicing QTLs (sQTLs) represent common genetic variants that influence gene expression levels and splicing patterns, respectively. To assess the effect of aseQC filtering on these signals, we used TensorQTL to map eQTLs and sQTLs in the ten GTEx tissues with the largest sample sizes and the highest number of aseQC-failed samples (Table S2, Fig. S11). QTL mapping was performed twice—once including all samples and once excluding those that failed aseQC. This comparison revealed no measurable difference in the number of eQTLs or sQTLs detected, indicating that removal of aseQC-failed samples does not disrupt the detection of cis-regulatory signals driven by common variants (Fig. S11A–B). This is expected, as aseQC-flagged samples represent only a small proportion of the total, comprising approximately 5.08% to 7.46% of samples in each of these GTEx tissues.

Transcriptome outlier detection is used to identify genes impacted by rare genetic variants, both in the general population and in rare disease cohorts [6, 8, 19-21]. In this context, the enrichment of transcriptional outlier genes for proximal rare, putatively disruptive variants serves as a key performance evaluation metric to benchmark outlier detection methods. We assessed changes in rare variant enrichment before and after excluding aseQC-failed samples across tissues in GTEx with over 25 samples. We quantified enrichment by calculating the odds ratio of a rare single nucleotide variant (minor allele frequency <0.1% in both GTEx and gnomAD) occurring within 10 kb upstream of the transcription start site in outlier versus non-outlier individuals, across gene expression, splicing, and ASE phenotypes across GTEx tissues with over 25 samples in-line with previous work [19]. Exclusion of aseQC-failed samples consistently increased rare variant enrichment for splicing and ASE outliers across tissues (Fig 3B-C). The most pronounced improvements were observed among ASE outliers with intermediate levels (p-value 0.01-0.001) of statistical significance (Fig. 3B, S12A–AW, S14A–AW). By contrast, gene expression outliers showed minimal or no improvement following aseQC filtering (Fig. 3B, S16A–AW). To further dissect the specificity of these effects, we stratified enrichment analyses by predicted variant consequence using annotations from the Variant Effect Predictor (VEP) [22]. Rare variant enrichment gains were consistently observed across high-impact variant classes—such as splice-site donors and acceptors and loss-of-function variants—for ASE and splicing outliers, while gene expression outliers again showed negligible change (Fig. 3C, S13A–AW, S15A–AW, S17A–AW). Together, these results demonstrate that while common variant-driven regulatory associations remain unaffected by filtering aseQC failed samples, aseQC substantially enhances the detection of rare regulatory variants particularly those influencing ASE and splicing signals.

## Discussion

Here, we introduce aseQC, a dedicated quality control framework for allele-specific expression (ASE) data. While traditional ASE analyses have relied on upstream quality control for genotyping and RNA-seq to ensure technical fidelity, methods that asses the quality of ASE data beyond broad genotype mismatches have been lacking. This gap has left ASE analyses vulnerable to latent technical artifacts, including sequencing errors, library preparation inconsistencies, and low-level contamination, which can subtly distort allelic balance without triggering existing QC filters. By modeling genome-wide allelic variability at the sample level, aseQC provides a quantitative measure of sample-wide variability in ASE data and enables systematic detection of distorted transcriptome samples that could otherwise compromise regulatory signal interpretation. In the GTEx dataset, aseQC identified hundreds of samples with inflated ASE variability that had passed all conventional DNA- and RNA-seq quality filters. These failures were enriched in specific tissues and clustered in time, most notably during a pronounced spike in September 2012, implicating temporal batch effects in RNA processing. Importantly, while gene expression outlier rates remained largely unaffected, ASE and splicing outliers were markedly inflated in these samples, reflecting the greater sensitivity of read-level transcriptomic features to technical noise. Removing aseQC-flagged samples substantially reduced these distortions and led to improved downstream analyses. The impact of aseQC-based filtering was particularly evident in rare variant enrichment analyses. We observed consistent and significant gains in enrichment for rare variants near transcription start sites when focusing on ASE and splicing outliers, indicating that poor ASE quality can obscure true regulatory effects. These improvements were largely absent in gene expression outliers, further supporting that more granular, read-level phenotypes are more susceptible to transcriptomic degradation. In contrast, cis-QTL mapping for common variants remained unaffected by aseQC filtering, even in tissues with high failure rates. This suggests that while large-scale QTL analyses are robust to modest levels of ASE degradation, rare variant analyses are more vulnerable and benefit directly from transcriptome-level quality control.

While aseQC presents a critical step to consider in transcriptome analysis pipelines, it is not without limitations. Its cohort-based nature makes it less applicable to single-subject analyses, and some sources of poor ASE quality remain difficult to trace with available metadata. Although batch date emerged as a strong predictor of aseQC score, traditional quality metrics like RIN or ischemic time showed limited explanatory power, suggesting the influence of unrecorded technical variables. Furthermore, a small number of previously annotated poor-quality samples in clinical rare muscular diseases were not flagged, indicating that additional axes of transcriptomic distortion may remain unmodeled. Nonetheless, aseQC offers a robust, interpretable, and computationally lightweight solution for improving the reliability of ASE analyses. As transcriptomic data continue to play an increasingly central role in rare disease diagnosis, variant prioritization, and regulatory genomics, it is critical that analytical pipelines distinguish biological signal from technical noise. By enabling sensitive detection of low-quality samples that compromise outlier detection and rare variant interpretation, aseQC strengthens the foundation of ASE-driven studies. Its performance across tissues, populations, and aggregation strategies supports its adoption as a standard quality control step for transcriptome-based analyses, ensuring that downstream inferences reflect genuine regulatory effects rather than artifact.

## Methods

### Statistical Model

Consider a cohort of J samples with gene aggregated ASE Data. Consider the reference counts (R) as an I-by-J matrix, where each element r_ij_ represents the reference count for gene i in sample j. Similarly, define the alternate counts (A) as an I-by-J matrix, with each element a_ij_ corresponding to the alternate count for the same gene-sample pair.

Therefore, matrix T representing the matrix of total counts for the corresponding genes/samples would be:

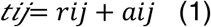

We capture the variability in allele-specific expression (ASE) across samples by utilizing a binomal logit-normal (BLN) distribution which is consistent with previous work [8]. Briefly, the reference counts are modeled in a Binomial distribution is:

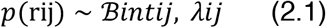

Where μij is the latent true reference ratio in sample j, for gene i, which is in-turn modeled via a logit-normal (LN) distribution as:

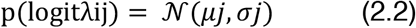

where *μ*_j_ and σ_j_ represent the mean and standard deviation of the logit-transformed reference ratios across all genes in sample j in log allelic fold-change unit [2].

We then use maximum likelihood estimation (MLE) to infer estimates for *μj, σj* for each sample. To ensure numerical stability of the model, we further incorporate a uniform component with a small weight, ε, resulting in:

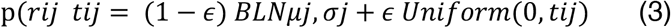

Hence, given the above framework in (3), the MLE estimates for *μj, σj* for sample_j_ would be obtained by maximizing:

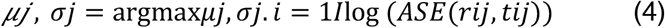

For each sample, the fitted *σ* has been termed the aseQC score, σ(BLN).

Within aseQC, only genes with total counts between 5 and 5000 are retained prior to fitting, and ε is set at 10^−3^. Maximum likelihood estimates (MLEs) (eqn. 4) are computed using the probability mass functions (PMFs) for the BLN distribution, provided by the previously published bln package [23]. The optimization is performed by minimizing the negative log-likelihood via the L-BFGS-B algorithm through the optim function in R.

### Outlier Detection

Given a cohort of ASE data, the aseQC first estimates the σ(BLN) for each sample. To determine a quality-threshold, a skew-adjusted boxplot is used which is implemented in the adjbox function in the robustBase package in R [11, 24]. The upper whisker of the boxplot fitted is termed σt, and samples with σ(BLN) > σt are considered to be aseQC outliers.

### Allele Specific Expression Data

#### Simulated Contamination data

Paired-end RNA-seq data were downloaded from the IGSR data portal for the GEUVADIS project, comprising 40 lymphoblastoid cell line (LCL) samples [3, 25, 26]. To simulate contamination, datasets were generated at 21 predefined levels (0%, 0.25%, 1%, 2.25%, …, 100%). For each contamination level, a specific fraction of reads corresponding to the non-contaminated proportion was sampled from the original source, while the remaining fraction was sampled from a designated contaminator sample, NA19159. For example, at a contamination level of 25%, 75% of the reads were sampled from the original source and 25% from the contaminator. Reads were sampled using seqtk (v1.0-r32) with a fixed random seed (−s100) to preserve paired-end reads. This process generated 840 FASTQ files across all contamination levels and samples. Genotype data for the contamination dataset were downloaded from the 1000 Genomes Project (https://ftp.1000genomes.ebi.ac.uk/vol1/ftp/data_collections/1000_genomes_project/release/20190312_biallelic_SNV_and_INDEL/) [26]. ASE data was subsequently produced using phASER (v1.0.0) with best-practice parameters, and genomic region blacklisting for GRCh38. Allelic counts were extracted from allelic_counts.txt files [10]. Subsequently, for each gene, the most expressed SNP (highest total count) was selected to represent gene-level ASE using bedtools (v2.27.1) [27]. Aggregated ASE data for both variant-level and haplotype-level were mapped to protein-coding genes using gencode v26 annotations. ASE Matrices derived from the synthetic contamination dataset can be accessed at https://github.com/PejLab/Contaminated_ASE. This dataset has also been archived on zenodo [28].

### GTEx

Sample-level variant ASE tables was downloaded from the protected data types for GTEx v8, and gene-level aggregations from variant-level ASE tables were performed by choosing the most expressed SNP for each gene consistent with previous work [4, 8]. Samples were identified using the “SAMPLE_ID” column in the available files and organized by tissue collected which were determined from publicly available sample attributes.

### DROP MAE QC Pipeline

The drop module (v.1.4.0) was installed using Bioconda [6]. Genotypes (in VCF file format) and RNA-seq (in BAM) were structured per input specifications. Subsequently, the DROP MAE QC module was executed using the published pipeline using snakemake (v.7.32.4) [29]. DROP MAE QC scores for samples were subsequently extracted from the file dna_rna_qc_matrix.Rds.

### Sample Attributes Correlated with ASE Quality in GTEx

Sample attributes predictive of σ(BLN) was investigated using a random forest model. All methods ultilized have been implemented in scikit-learn (v.1.3.2) in python (v.3.8.3) [30]. Publicly available sample attributes for GTEx v8 were downloaded from the GTEx portal [4]. Attributes were classified into three broad classes: “RNA-sequencing” (derived from RNASeqC), “temporal” (connected to the date of sequencing), and “other/pre-sequencing” (encompassing information known prior to sequencing) (Table S4). Among features in the “other/pre-sequencing” category, an Iterative imputer was used to impute values for ischemic times (SMTSISCH), autolysis scores (SMATSSCR), and PaxGene times (SMTSPAX) if values were missing. One-hot encoding was used to convert tissue-names to vectors. Following this, all columns with non-numeric values were dropped (Table S4). Four broad classes of models were then fitted to evaluate the impact of various classes of attributes, namely “without date without RNA-seq”, “without date with RNA-seq”, “with date without RNA-seq”, “with date with RNA-seq”. These models include/exclude features from the feature classes. The model “without date without RNA-seq”, includes features from just the “other/pre-sequencing” class and serves as a base-model to evaluate performance. Following this, the feature matrix (X) is constructed from features belonging to the respective attributes with non-null values, and the predictor variable (y) is σ(BLN). The train_test_split function from scikit-learn is used to set a training/test set using a test_size=0.2 (20% of the total data size) [30]. A RandomForestRegressor was trained, and hyperparameters were optimized through a grid search with 3-fold cross validation (Fig S6A). The hyperparameters explored included: n_estimators: 100, 200; max_depth: None, 10, 20; min_samples_split: 2, 5; min_samples_leaf: 1, 2. The performance metric for model selection was the negative mean squared error. Following this, predictions on the test set, σ(BLN)_predicted_ were generated, and then stratified by tissue. For each tissue within the test-set, the upper-whisker of the skew-adjusted boxplot (robuseBase::adjboxstats) was then generated based on the predictions, and classifications á la aseQC were made (i.e. fail if σ(BLN)_predicted_ > σ_t_predicted_) [11, 24]. These predicted classifications were then compared with the actual labels which were generated from aseQC. The performance of the predictions are then assessed with a Precision, Recall, F1-score and Mathews correlation coefficient (MCC (ϕ); r-score) as implemented in sklearn.metrics [30].

### Gene Expression PCA

Gene expression data (were downloaded from the GTEx portal. Protein-coding genes were defined based on gencode v26 annotations and retained if their median expression across individuals exceeded 100 counts. Raw counts were log_10_transformed after adding a pseudo-count of 10 to stabilize variance and mitigate the influence of low-expression genes. Principal component analysis (PCA) was then performed using the scikit-learn implementation (v.1.3.2) in Python (v. 3.8.3), with the first four principal components retained for downstream analysis, collectively explaining over 50% of the total variance (Table S5, [30]). To quantify global expression deviation, samples were stratified by tissue, and a centroid was computed in the resulting four-dimensional PCA space for each tissue. The Euclidean distance from this centroid in 4-dimensional space was then calculated for each sample, allowing us to assess whether aseQC-failed samples were systematically displaced from the tissue-specific expression norm.

### Gene Expression Outlier Detection

Gene expression data (read counts) were downloaded from the GTEx portal and then stratified by tissue [4]. For each tissue with over 25 samples, protein-coding genes (as defined in gencode v26) with a median greater than 100 counts across all samples were retained for analysis. The OutSingle pipeline was subsequently run for each tissue for expression outlier detection using Python (v.3.8.3) [17]. Briefly, gene expression data for each tissue was structured to input specifications, and the fast_zscore_estimation.py, and optht_svd_zs.py scripts were used to obtain scores for each sample. OutSingle z-scores were converted to two-tailed p-values using the stats.norm.cdf function from SciPy (v.1.12.0) [31]. These p-values were then corrected for false discovery rate (FDR) using the Benjamini-Hochberg procedure, implemented via multitest from statsmodels.stats (v. 0.13.2) with a significance cutoff of 0.05 [32].

### Splicing Outlier Detection

RNA-seq BAM files were retrieved from the GTEx portal’s protected-access data and stratified by tissue. For each tissue with more than 25 samples, FRASER2 was run using R (v4.3.3) installed via Bioconda [18]. Data preprocessing included read counting and the calculation of Percent-Spliced-In (PSI) values. Expression filtering was applied to retain splicing events with sufficient expression in at least one sample (minimum expression threshold = 20 reads, 75th quantile cutoff = 10 reads). The FRASER2 model was used to identify splicing outliers, employing Jaccard-based quantile normalization, using an autoencoder dimension of (q = 3). Genomic ranges were annotated using the TxDb.Hsapiens.UCSC.hg38.knownGene database, with gene-level annotations derived from org.Hs.eg.db [33, 34]. Gene-level p-adjusted values were calculated and FDR correction was performed on these gene-level aggregated p-values using a Benjamini-Hochberg procedure, with a significance cutoff of 0.05.

### ASE Outlier Detection

Outlier analysis was conducted using ANEVADOT package (v.0.1.1) in R (v.4.0.0). Gene-aggregated, variant-level ASE data were obtained for each sample and stratified by tissue, focusing on tissues with more than 25 samples [8]. V^G^s derived from GTEx v8 data for the corresponding tissues, were downloaded from the publicly accessible GitHub repository (https://github.com/PejLab/ANEVA-DOT_reference_datasets/tree/master/Reference_Vg_Estimates). The ANEVADOT_test function was applied to each sample using default parameters and tissue appropriate V^G^s. For each sample, the “adj.pval” value represents the Benjamini-Hochberg false discovery rate (FDR)-adjusted p-value subsetted on protein-coding and lnc-RNA genes as defined in gencode v26. A significance threshold of 0.05 was used to identify outliers.

### Expression and Splicing QTL Analysis

The ten GTEx tissues with the highest proportion of samples rejected by aseQC score were chosen for eQTL and sQTL analysis (Table S3). Gene expression and alternative splicing were quantified for each tissue using Pantry as previously described [35]. Pantry was then used to map cis-eQTLs and cis-sQTLs using tensorQTL, using the default Pantry covariates of 20 principal components from the RNA phenotype table and 5 principal components from the genotypes [36]. sQTL mapping was run with the group setting to handle multiple phenotypes per gene. For both eQTL and sQTL mapping, tensorQTL was run in cis mode and then cis_independent mode to generate zero or more conditionally independent QTLs per gene. This procedure was run with the full sample set and again with the aseQC-filtered sample set per tissue.

### Rare Variant Enrichment

Single nucleotide variants (SNVs) with a minor allele frequency (MAF) <0.1% (as reported in GTEx and gnomAD v2.0.2) were identified from GTEx genotype data, and their functional consequences were predicted using the Variant Effect Predictor (VEP) for each individual in-line with our previous effort [19, 22]. Variants were assigned to genes if located within a ±10 kb window of the gene boundary. Relative risk calculations for rare variant enrichment were performed using the ExOutBench framework (v1.0.0) in R (v4.0.0), following methodologies consistent with prior published enrichment analyses [19]. Briefly, relative risk was estimated by selecting genes with significant outlier scores in at least one individual for transcriptional phenotypes i.e. (gene expression, splicing, or allele-specific expression (ASE)). The relative risk was quantified as the odds ratio of an outlier individual carrying the variant compared to a non-outlier individual for the respective phenotype. Enrichment scores were further refined by stratifying results across varying outlier significance thresholds and by VEP annotation categories using the enrichment_by_significance, and enrichment_by_annotations functions with default settings in ExOutBench [22].

## Author Contributions

KG led data processing, analysis, interpretation, and manuscript preparation. ES contributed to the development of the statistical framework. DM performed QTL mapping and assisted in result interpretation. AT provided analytical support and contributed to data interpretation. PM conceived the study, supervised the analyses, and edited the manuscript. All authors have reviewed and approved the final manuscript.

## Data and Code Availability

The allele-specific expression (ASE) data analyzed for this study for GTEx are available to authorized users through dbGaP under accession no. phs000424.v8 and on the GTEx portal (https://gtexportal.org/). The statistical model and code underlying aseQC are available as an R package at https://github.com/PejLab/aseqc_repo/. The method is also available within a container at https://hub.docker.com/repository/docker/krganapathy/ase_outlier_tools/. Reference ASE data from the synthetic contaminated dataset is available at https://github.com/PejLab/Contaminated_ASE. All data and code produced in this study has been made publicly available on zenodo [12, 13, 28].

## Acknowledgements

We would like to sincerely thank Robert Vogel, Douglas Evans, Chunlei Wu, Andrew Su, Nathan Wineinger and Tuuli Lappalanien for helpful discussions on this project. We thank JC Ducom and Lisa Dong from the Scripps Research HPC Team for computational infrastructure support. We thank the GTEx donors for their contributions to science, the GTEx Laboratory, Data Analysis, and Coordinating Center (LDACC), and the GTEx analysis working group (AWG) for their work in generating the resource.

