## Supplementary Methods for "Allele Specific Expression Quality Control Fills Critical Gap in Transcriptome Assisted Rare Variant Interpretation": Supp_Text_Fixed_Ref_June_7.pdf


### Supplementary Images

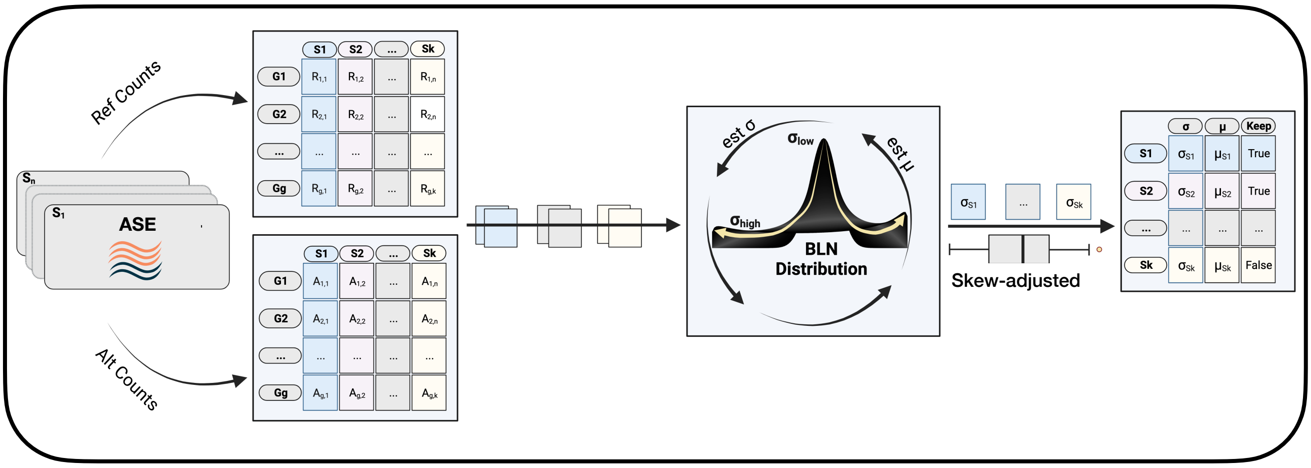


**Figure S1: Workflow to Quantify Quality in Sample-Level ASE Data Using BLN Distribution**
Schematic representation of the aseQC pipeline for cohorts, detailing the calculation of σ for each sample with the application of a skew-adjusted boxplot to establish quality thresholds [1]

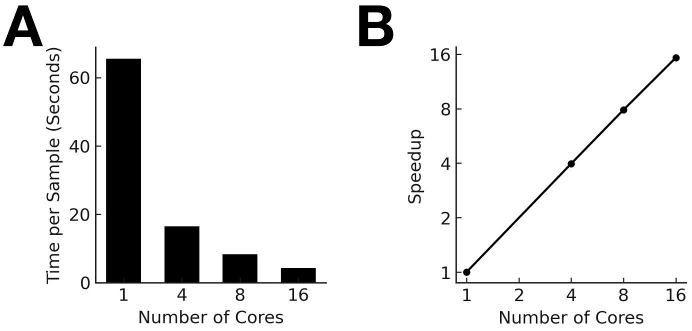


**Figure S2: Runtime Evaluation of aseQC**

**(A)** Per-sample runtimes for 64 identical ASE samples, showing the effect of varying the number of cores on aseQC fit performance. **(B)** Speed-up achieved by aseQC relative to single-core performance as the number of cores increases.

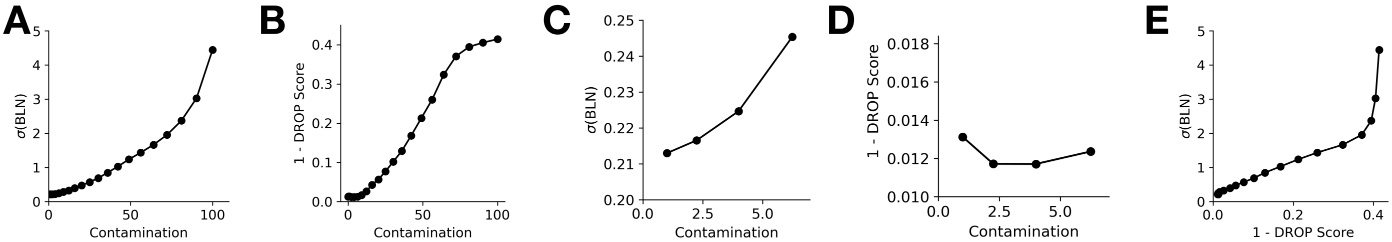


**Figure S3: Comparing Detection of Sample Mismatches by aseQC and DROP-MAE for Sample HG00139**
**(A)** aseQC shows a monotonically increasing σ(BLN) with rising contamination. **(B)** DROP-MAE QC exhibits a sigmoid response curve, plateauing at higher contamination levels. **(C)** aseQC is more responsive particularly at higher contamination levels.

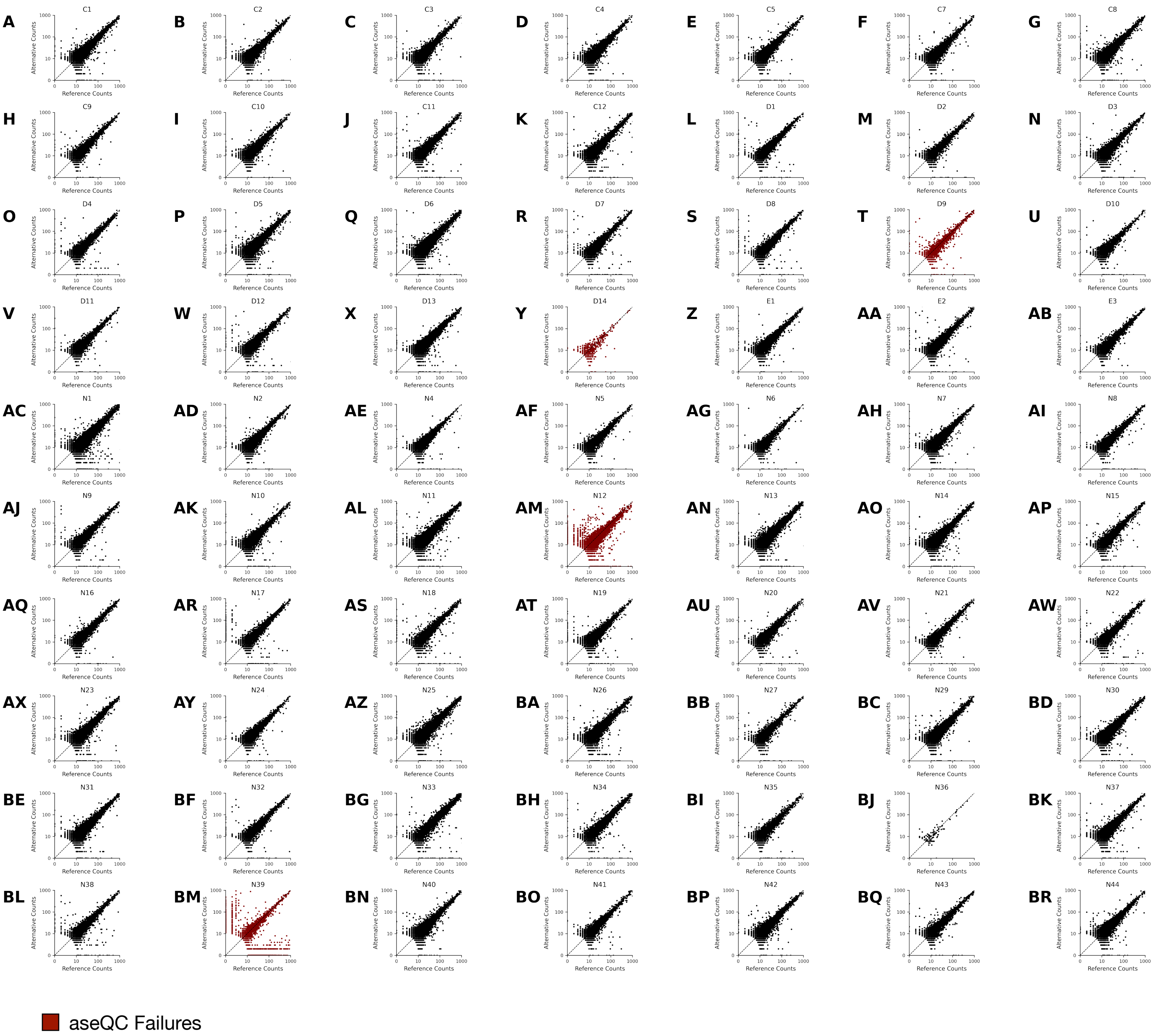


**Figure S4: Applying aseQC to a Cohort of 70 Muscular Dystrophy Cases**

Applying aseQC identifies 4 samples (in red) that failed the aseQC test. 2 of these samples (panels BM, AM) are known previously to be of low quality.

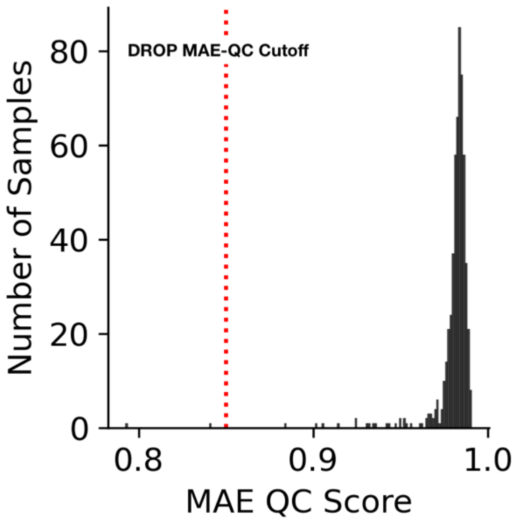


**Figure S5: Comparison of DROP MAE QC scores among Samples Failing aseQC**
Histogram depicting DROP MAE-QC scores for 563 samples that failed aseQC. The dotted red line represents the DROP MAE-QC passing threshold. Notably, all but two aseQC-failing samples meet the DROP MAE-QC criteria suggesting no genotype-RNA mismatches.

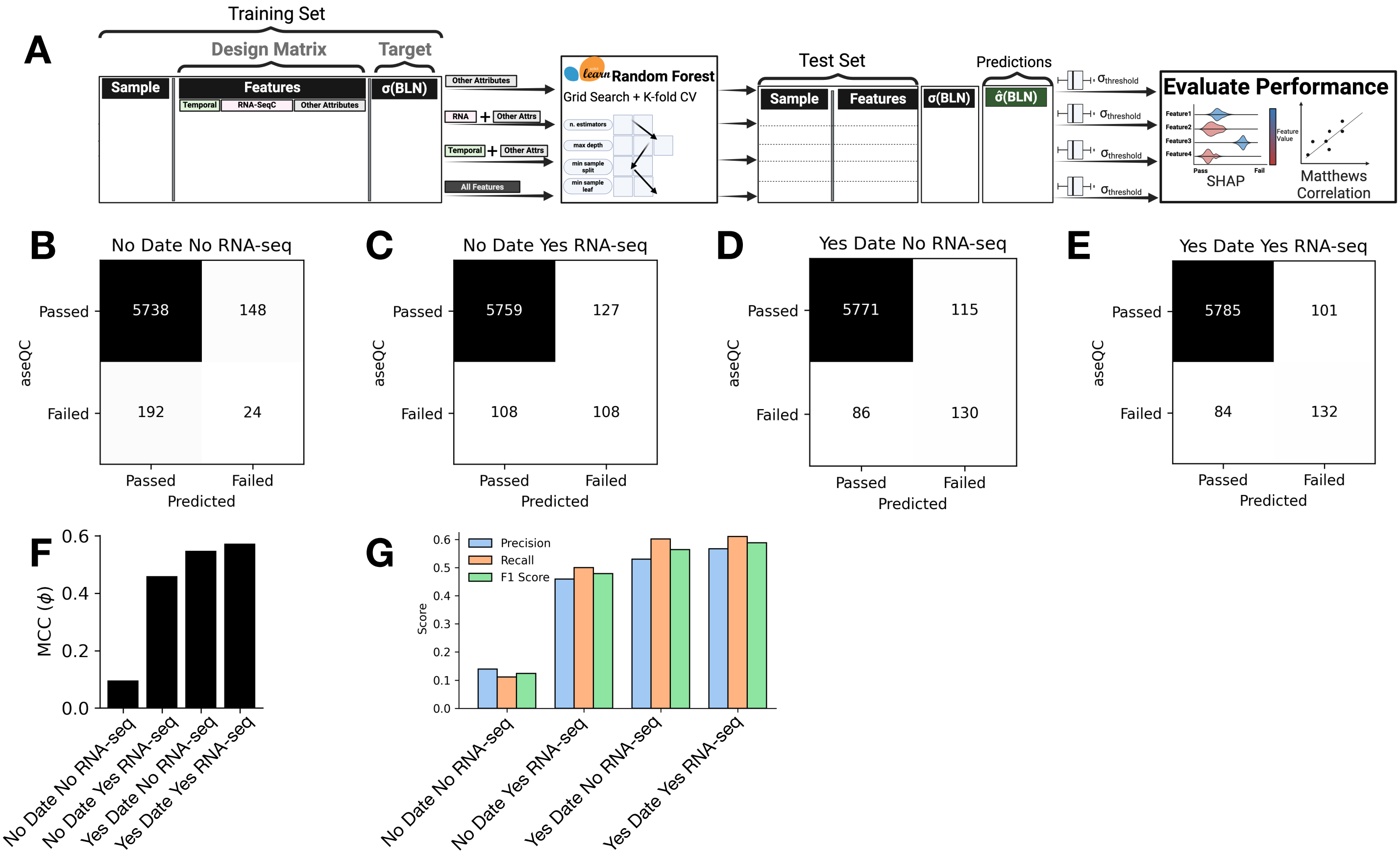


**Figure S6: Evaluation of Random Forest Models for Predicting aseQC score from Sample Attributes**
**(A)** Schematic workflow illustrating the Random Forest model fitting process to identify attributes linked to ASE sample quality [2]. **(B-E)** Confusion matrices depicting prediction performance on the test set across four models with different feature sets. **(F)** Matthew's Correlation Coefficient (MCC ($\phi$); r-score) for the test set across the four models. **(G)** Precision, Recall, and F1 scores for all four models on the test set.

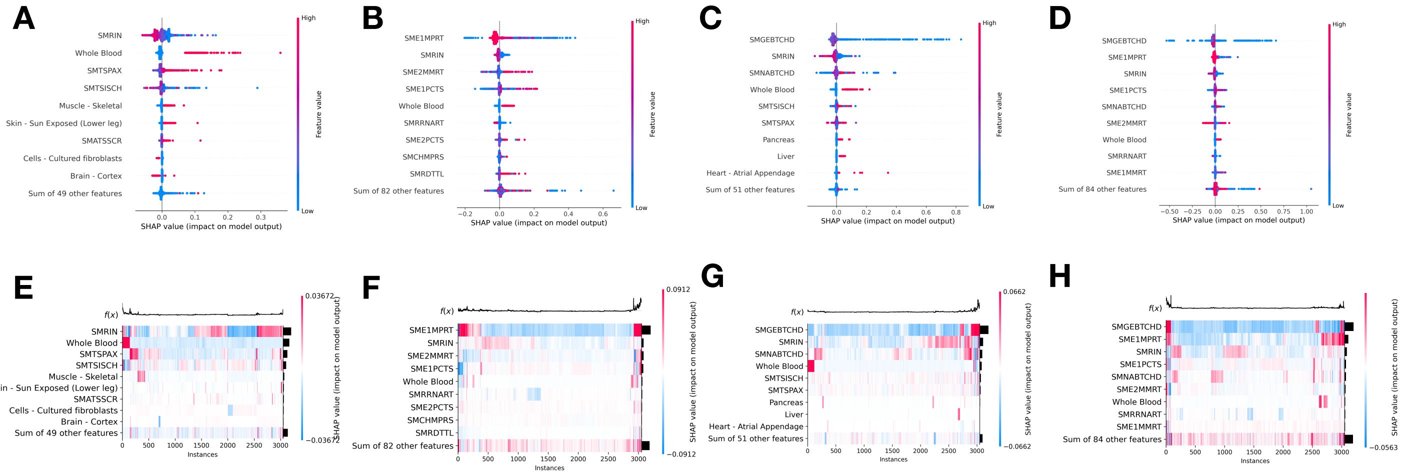


**Figure S7: SHAP Plots Highlighting Features Contributing to aseQC Score Prediction**
**(A-D)** Beeswarm plots illustrating how feature values influence SHAP scores, driving predictions toward lower values (aseQC pass) or higher values (aseQC failure). **(E-H)** Heatmaps depicting the interactions among features, showcasing their combined impact on predicting aseQC passes or failures across all four models.

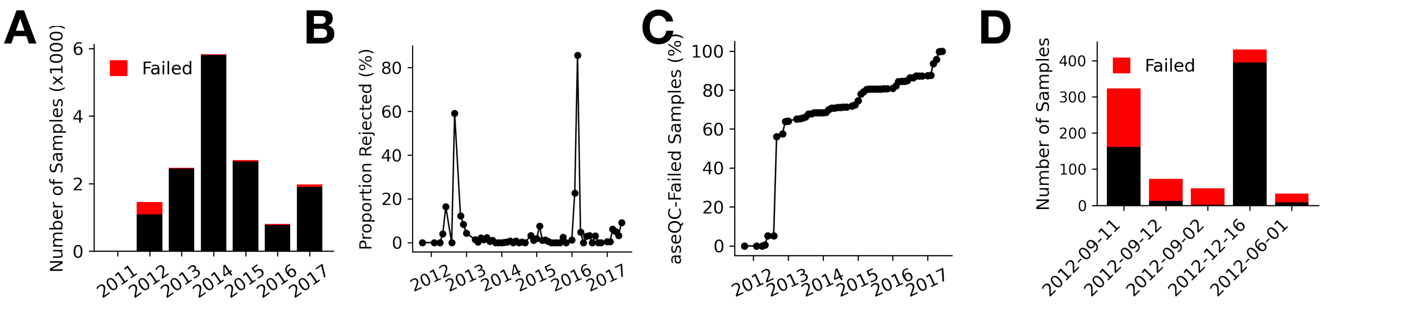


**Figure S8: aseQC Failures by Sample Processing Date**
**(A)** Number of aseQC Failures aggregated per year **(B)** Proportion of aseQC sample failures aggregated by month. **(C)** Cumulative distribution of the proportion of total samples failed. **(D)** Top 5 days with the highest number of aseQC failures.

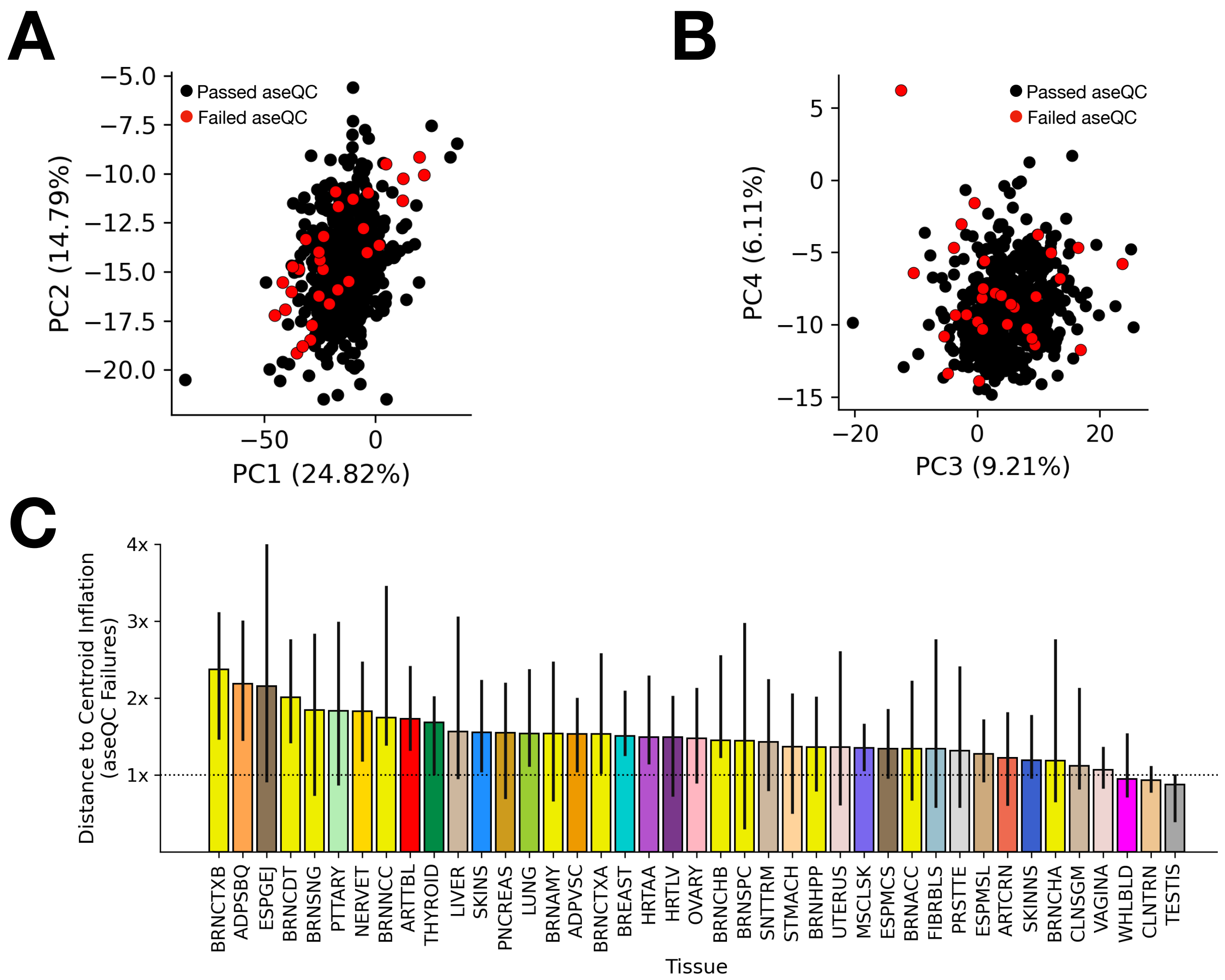


**Figure S9: PCA-Based Assessment of Global Gene Expression Deviation in aseQC-Failed Samples Across GTEx Tissues.**

**(A)** PC1 vs PC2 and **(B)** PC3 vs PC4 projections of log-transformed gene expression profiles for Adipose Subcutaneous tissue, with samples colored by aseQC status (black = passing, red = failing). **(C)** Fold-change in median Euclidean distance to the tissue-specific centroid, defined as the average position of all samples in four-dimensional PCA space, comparing aseQC-failed to passing samples across GTEx tissues with >25 samples and at least 5 failures. Error bars represent 95% bootstrap confidence intervals.

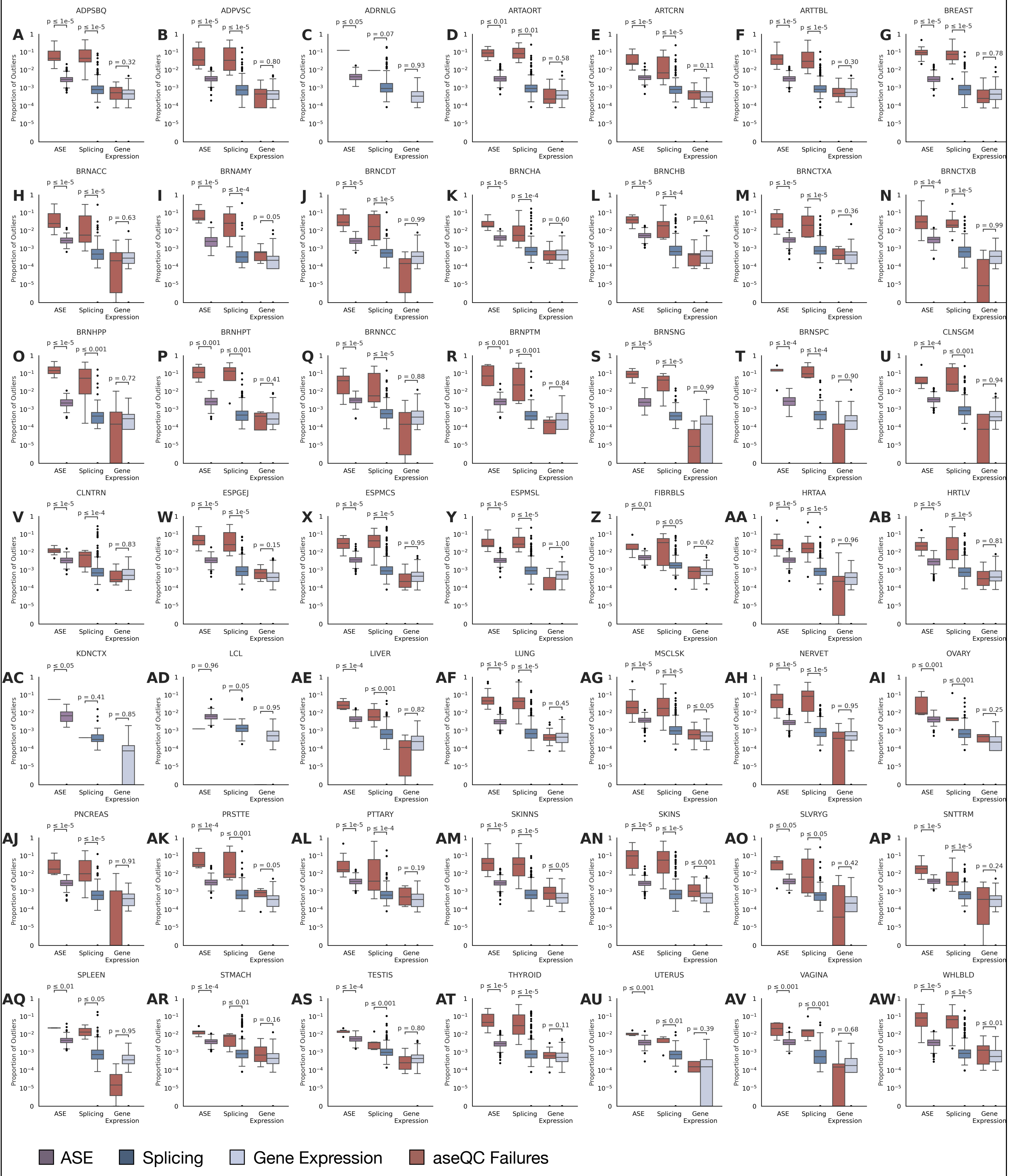


**Figure S10: Proportion of outlying genes identified in samples categorized as passed or failed by aseQC.** **(A-AW)** Outlying genes were detected using ANEVA-DOT for allele-specific expression (ASE), FRASER2 for splicing outliers, and OutSingle for gene expression outliers for all GTEx tissues with over 25 samples.

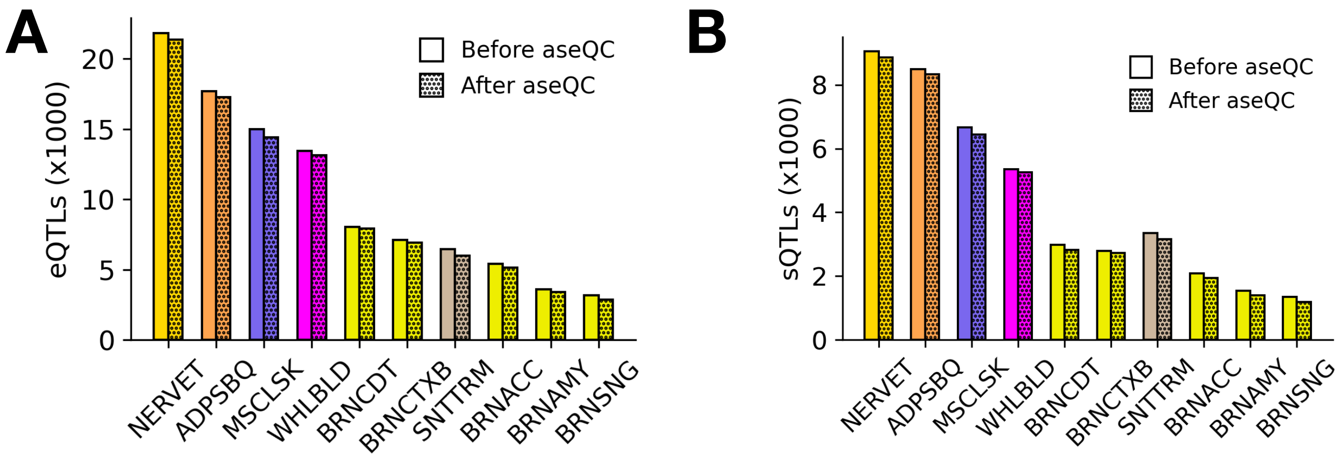


**Figure S11: Effect of aseQC-based filtering on cis-QTL discovery in GTEx tissues.**
**(A)** Number of significant eQTLs and **(B)** sQTLs identified before and after excluding aseQC-failed samples, shown for the top-ten GTEx tissues (n > 25) with most proportion of aseQC failures

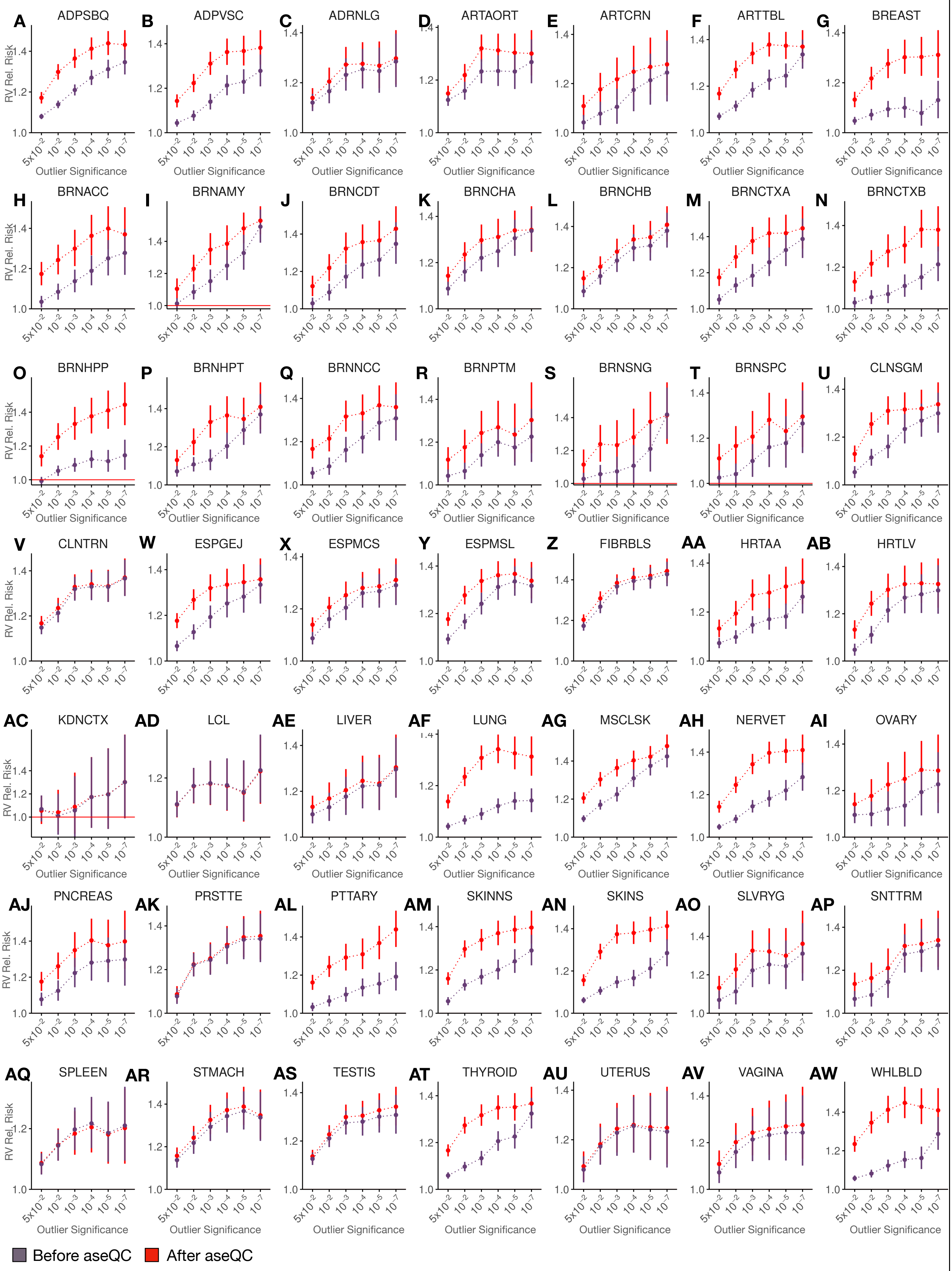


**Figure S12: Changes to ASE Rare Variant Enrichment by Outlier Significance Post-aseQC Filtering (A-AW)** Comparison of changes to rare variant enrichment stratified by DOT outlier significance after removing aseQC failures in 49 GTEx tissues.

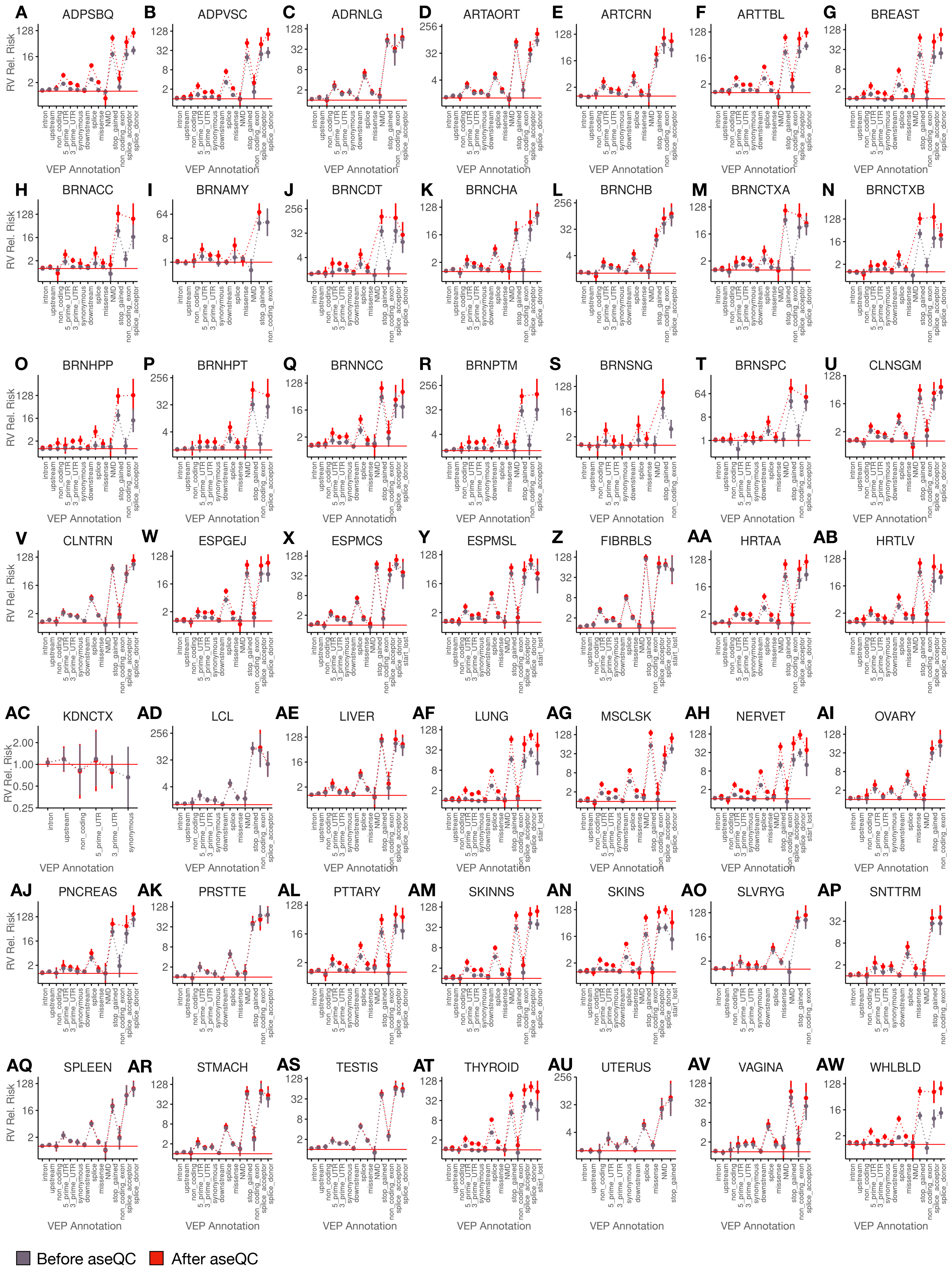

**Figure S13: Changes to ASE Rare Variant Enrichment by VEP Consequence Post-aseQC Filtering**
**(A-AW)** Comparison of changes to rare variant enrichment among DOT outliers (FDR qval <= 0.05) stratified by VEP consequence in variant- and haplotype-aggregated ASE data in 49 GTEx tissues.

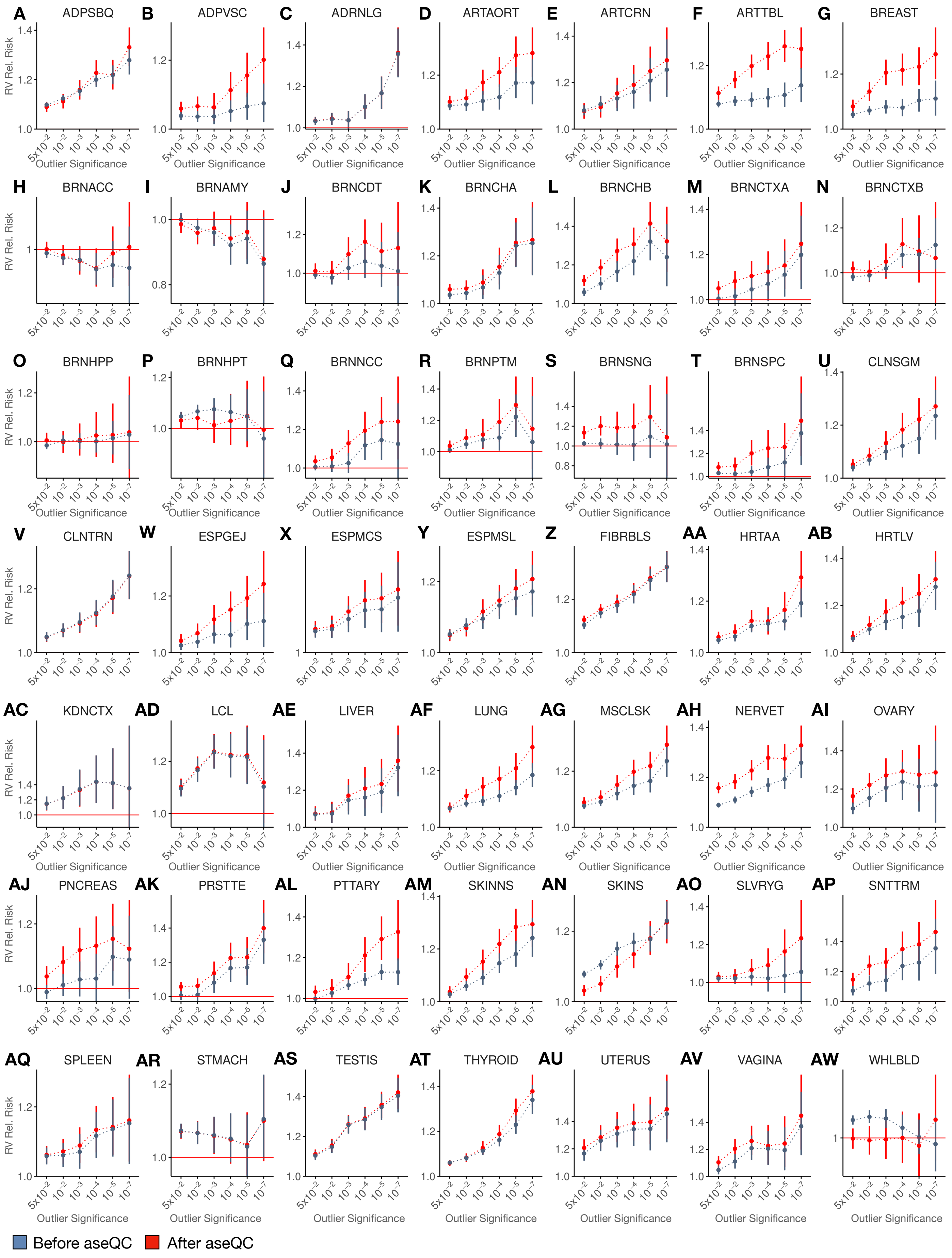


**Figure S14: Changes to Splicing Rare Variant Enrichment by Outlier Significance Post-aseQC Filtering (A-AW)** Comparison of changes to rare variant enrichment stratified by FRASER2 outlier significance after removing aseQC failures in 49 GTEx tissues.

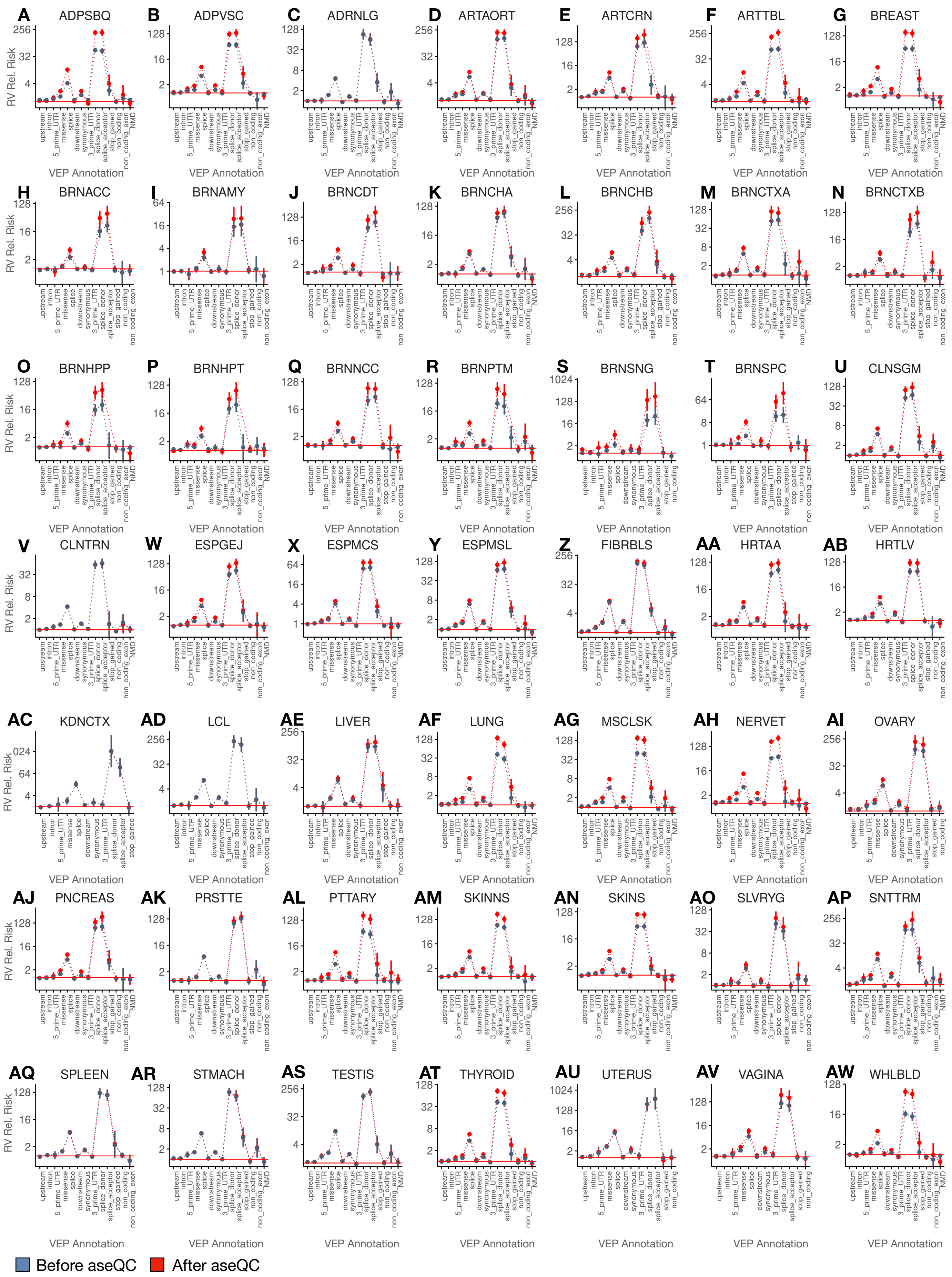


**Figure S15: Changes to Splicing Rare Variant Enrichment by VEP Consequence Post-aseQC Filtering (A-AW)** Comparison of changes to rare variant enrichment among FRASER2 outliers (FDR qval <= 0.05) stratified by VEP consequence in variant- and haplotype-aggregated ASE data in 49 GTEx tissues.

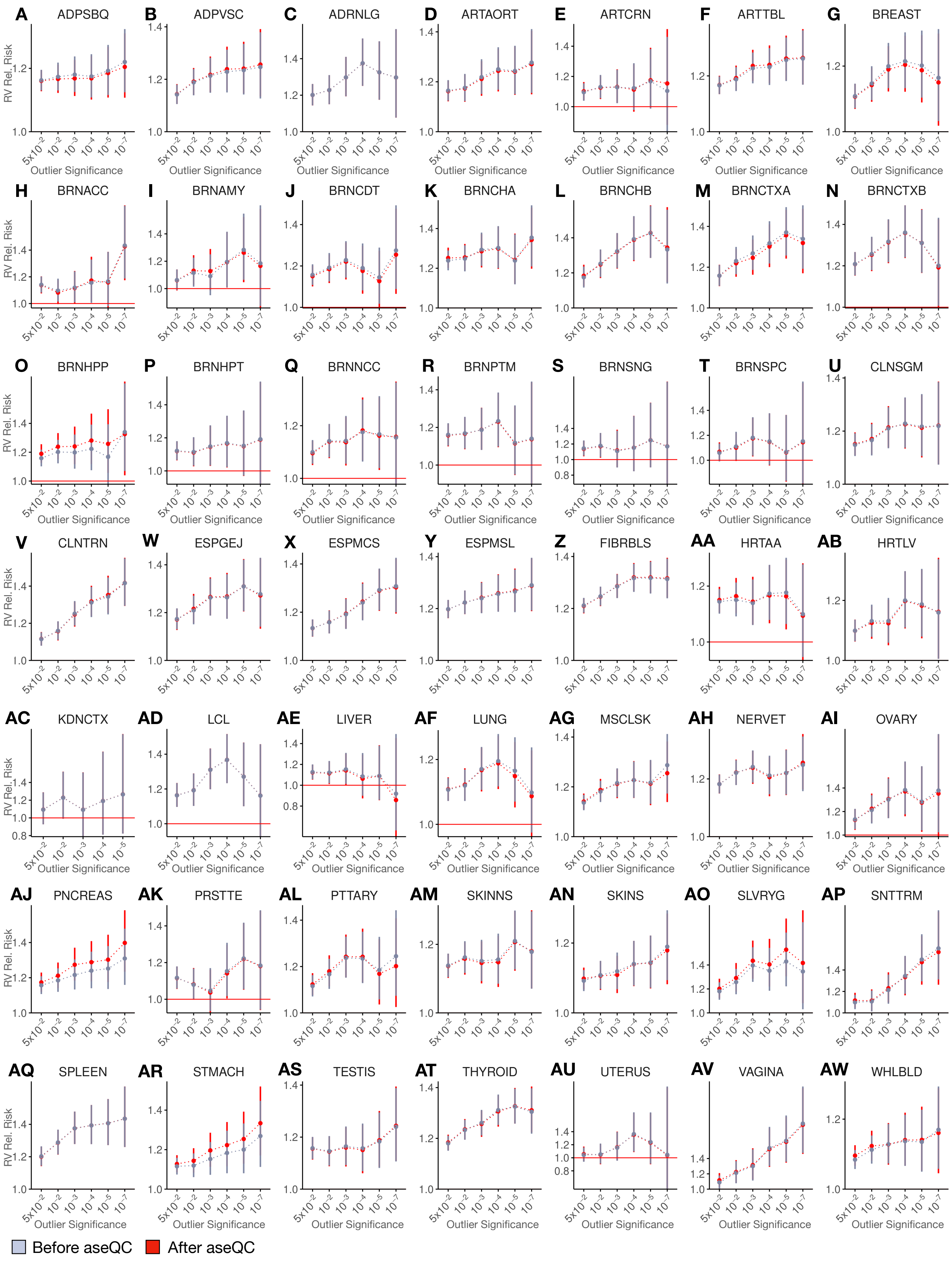


**Figure S16: Changes to Gene Expression Rare Variant Enrichment by Outlier Significance Post-aseQC Filtering**

**(A-AW)** Comparison of changes to rare variant enrichment stratified by OutSingle outlier significance after removing aseQC failures in 49 GTEx tissues.

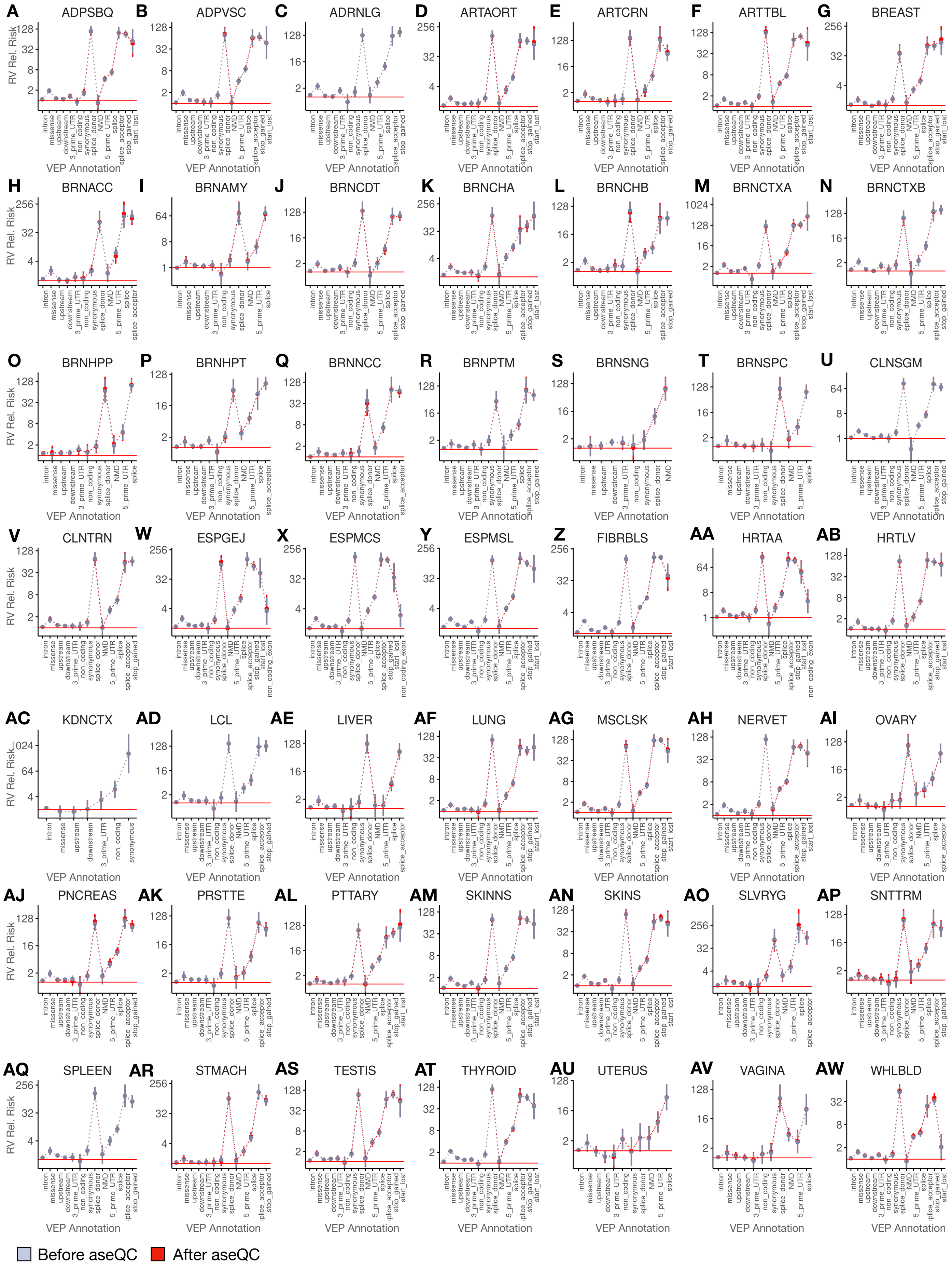


**Figure S17: Changes to Gene Expression Rare Variant Enrichment by VEP Consequence Post-aseQC Filtering**

**(A-AW)** Comparison of changes to rare variant enrichment among OutSingle outliers (FDR qval <= 0.05) stratified by VEP consequence in variant- and haplotype-aggregated ASE data in 49 GTEx tissues.
